# Early life stress affects the transcription and chromatin accessibility of spermatogonial cells

**DOI:** 10.64898/2026.08.20.745941

**Authors:** Rodrigo G. Arzate-Mejía, Theresa Schöpp, Kerem Uzel, Irina Lazar-Contes, Isabelle M. Mansuy

## Abstract

Adversity in early life has lasting effects on the physiology and behavior of exposed individuals and their descendants. In mice, early-life stress alters the RNA content of adult sperm, and this RNA is sufficient to transmit some of the effects to the offspring who were never exposed. However, sperm cells are not yet formed during the early postnatal window in which the exposure occurs. Spermatogonial cells (SPGs), which give rise to them, are present at that time, but whether they respond to the exposure and maintain a molecular signature of it into adulthood is unknown. Here we show that early-life stress alters both the transcriptome and the chromatin accessibility of mouse SPGs, and that a molecular signature of the exposure remains detectable in adulthood. One day after exposure ended, the transcriptional response was extensive, with proliferation and nucleosome-organization programs coordinately up-regulated. In adulthood, the transcriptional response was modest and dominated by coordinately down-regulated gene programs. Single-cell profiling of the whole testis localized the adult response to spermatogonial stem cells (SSCs) and to genes involved in spermatogenesis. At the chromatin level, accessibility shifted one day after exposure at binding motifs for signal-responsive transcription factor families, and in adulthood at a different set of families, in both cases at primed enhancers. These data demonstrate that SPGs respond to an early postnatal environmental exposure and identify them as a candidate origin of the molecular changes later found in adult sperm.

## INTRODUCTION

Adverse experiences in early life are common in humans and increase the risk of psychiatric and metabolic disease across the lifespan^1,2^. In mice, exposure to chronic stress during the first two weeks of postnatal life induces lasting effects on physiology, cognition and behavior in the affected adult males ^3–5^, and comparable effects are observed in their offspring^6^, despite never being exposed to the stressor themselves.

Environmental exposures can leave a molecular trace in the male germline. Paternal diet, gut microbiota perturbation, and traumatic stress each alter the RNA content of mature sperm^6–8^, and for early-life stress this altered RNA cargo is sufficient for intergenerational transmission of some of the observed phenotypes^6^. The sperm carrying these changes, however, did not exist at the time of exposure, since the first haploid spermatids appear only around postnatal day (PND) 20, and the first wave of spermatogenesis is not complete until PND30–35^9,10^. The information must therefore have been registered in a germ cell that was present during the exposure, that gives rise to sperm, and that retains the change until sperm are formed.

The germ cells available to meet these requirements are set by the sequence of postnatal testicular development. Prospermatogonia are exhausted by PND5, and distinct spermatogonial subtypes can be distinguished from PND6–8^10,11^. Spermatogonial stem cells (SSCs) either self-renew or generate differentiating spermatogonia, which form the primary and secondary spermatocytes that appear around PND10–12; these enter meiosis and give rise to haploid spermatids that develop into spermatozoa^12–14^. Therefore, during the first two postnatal weeks the testis contains SPGs and early meiotic cells, but no post-meiotic germ cells. Of these, SPGs are among the most abundant^10^, they give rise to every subsequent spermatogenic stage^15,16^, and they alone contain a self-renewing stem cell population that persists throughout reproductive life^17^. Although SPGs satisfy these requirements and their development and self-renewal are well characterized, little is known about whether they respond to early-life environmental exposures or whether such responses are sustained.

## RESULTS

### Early-life stress alters the spermatogonial cell transcriptome

To study the short- and long-term molecular effects of early-life stress (ELS) on mouse SPGs, we used a model of unpredictable maternal separation combined with unpredictable maternal stress (MSUS), an established paradigm with intergenerational effects^5,6^. F1 pups underwent MSUS from PND 1 to 14 (Supplementary Fig. 1a). Exposed pups had lower body weight than controls at PND7 and PND14, indicating a physiological response to the paradigm during the exposure itself (Supplementary Fig. 1b). In adulthood, exposed males had lower body weight and spent a greater proportion of time in the open arms of an elevated plus maze (Supplementary Fig. 1b,c), confirming the long-term physiological and behavioral effects of the MSUS paradigm^6,18^.

Then, we assessed the short-term transcriptional effects of ELS on SPGs one day after exposure ended (PND15). SPGs (CD9⁺CD45⁻CD51⁻c-KIT⁻) were FACS-enriched from control and ELS F1 males and profiled individually by RNA-seq. The sorted fractions expressed spermatogonial and germ cell markers with minimal somatic signal (Supplementary Fig. 2j–l). Deconvolution analysis estimated that SPGs accounted for over 97% of the cells in each RNA-seq library, showing no difference in composition between control and ELS groups (Supplementary Fig. 2j–l). Principal component analysis (PCA) distinguished the two groups, indicating that ELS influences the PND15 SPG transcriptome (Fig. 1a). At the gene level, 219 genes were differentially expressed (FDR < 0.05; median absolute log2FC 0.235; Fig. 1b), with 132 upregulated and 87 downregulated. Upregulated genes included chromatin regulators (*Chd1*, *Ncapd2*, *Lmnb1*), nucleosome components (*H2ac1*, *H2bc1*, *H3f4*, *H1f2*, *H1f1*, *H4c11*), and spermatogonial factors associated with spermatogonial stem cell (SSC) identity and self-renewal (*Bcl6b*, *Spry4*, *Epha2*, *Foxc2*, *Etv5*, *Dusp6*, *Dusp9*, *Etv4*, *Gfra1*). Downregulated genes included transcriptional and epigenetic regulators (*Pax7*, *Zim2*, *Kcnq1ot1*, *Tet3*) and members of the piRNA pathway (*Piwil4*, *Tdrd9*, *Tdrd5*) (Fig. 1c).

**Fig. 1.**
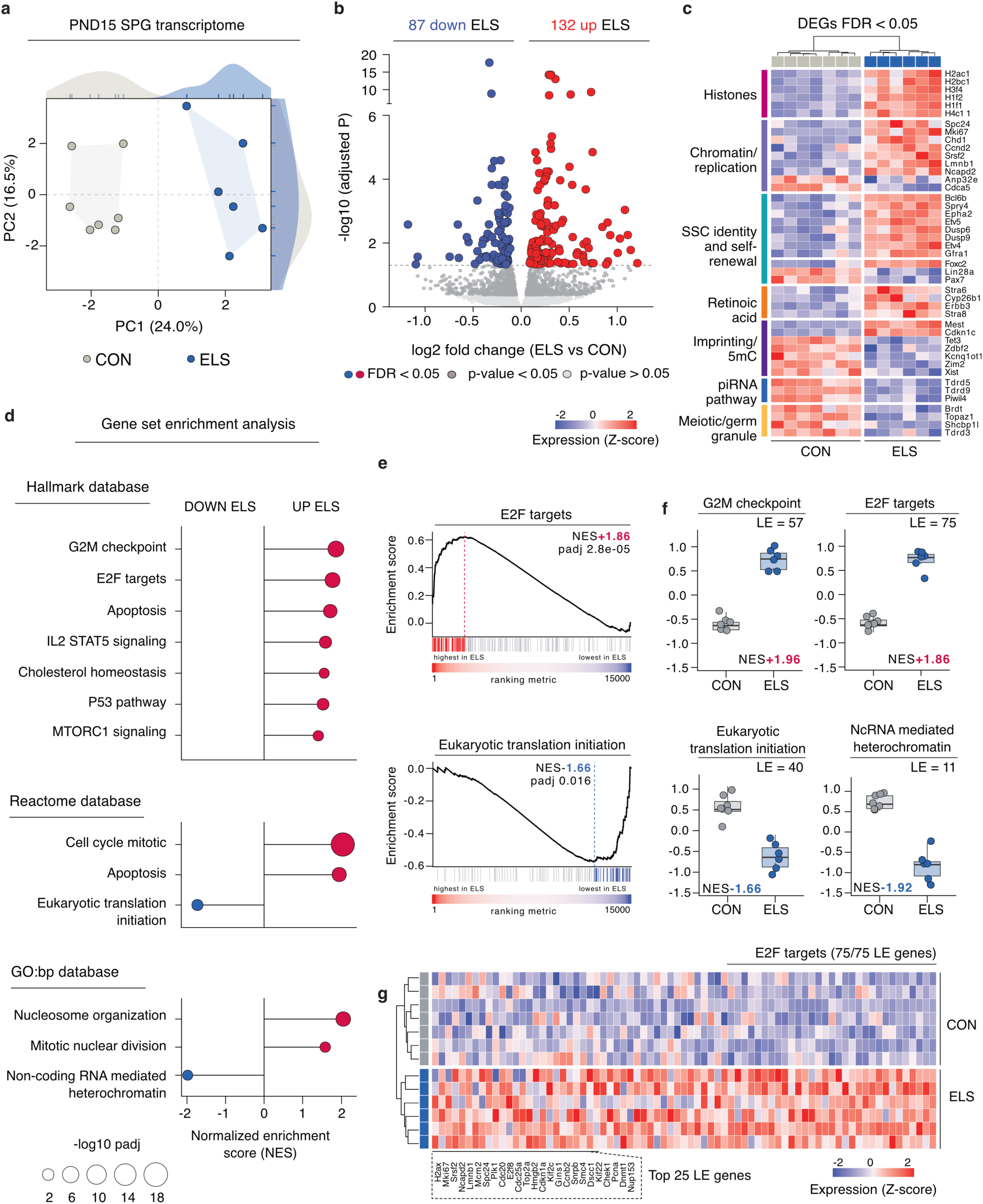
Early-life stress alters the spermatogonial transcriptome at PND15. **a**, Principal component analysis of the SV-corrected transcriptome from spermatogonia (SPG) from control (CON) and early-life stress (ELS) F1 males at PND15. Axes show the variance explained by each component. *P* value from a two-sided Wilcoxon rank-sum test on PC1 (CON *n* = 7; ELS *n* = 6 males; *P* = 0.0012). **b**, Volcano plot of differential gene expression between ELS and control SPGs. Color marks genes at FDR < 0.05 (red, higher in ELS; blue, lower), nominal *P* < 0.05 (mid grey), and *P* > 0.05 (light grey); counts above the panel give genes in each direction. *P*-values were obtained from the edgeR likelihood ratio test, adjusted for multiple testing (Benjamini–Hochberg) (CON *n* = 7; ELS *n* = 6 males). **c**, Heatmap of 44 of the 219 differentially expressed genes (FDR < 0.05), selected to represent seven functional categories. Each gene shown is individually significant, and the six histone genes are representatives of 26 significant replication-dependent histone genes. Columns are individual males (CON *n* = 7; ELS *n* = 6), clustered on gene-wise *z*-scores (dendrogram above); rows are grouped by category. **d**, Gene set enrichment analysis of the Hallmark, Reactome and GO:BP collections. Point position, normalized enrichment score (NES); point area, -log10 adjusted *P*; color, direction of change. Sets shown are non-redundant representatives of those at FDR < 0.05 (Hallmark 7 of 13, Reactome 3 of 109, GO: BP 3 of 19); Cell cycle mitotic is a Reactome ancestry domain covering 24 nested pathways, and Apoptosis appears in the Hallmark and Reactome blocks as two distinct sets. **e**, Running enrichment score for E2F targets (top) and eukaryotic translation initiation (bottom) across genes ranked by the signed likelihood-ratio statistic. Vertical lines mark gene set members; colored lines are the leading-edge (LE) genes for that set; the strip below shows the ranking metric from highest to lowest in ELS. **f**, LE module score per male for four enriched sets, computed on the SV-corrected assay as the mean per-gene *z*-score of the leading-edge genes of each set. Box, interquartile range with median line; whiskers, 1.5× interquartile range; points, individual males (CON *n* = 7; ELS *n* = 6). Scores are not sign-adjusted, so sets lower in ELS sit below zero. **g**, Heatmap of all 75 E2F-target LE genes, *z*-scored per gene. Rows are individual males, clustered (dendrogram at left) and annotated by group at right; columns are the leading-edge genes in ranking order, the 25 highest-ranked labeled.

Next, we applied gene set enrichment analysis (GSEA) to identify coordinated shifts in transcriptional programs across gene sets associated with specific biological processes. Enrichment analysis revealed upregulation of gene sets involved in cell cycle and chromatin regulation, including G2M checkpoint (NES +1.96, p-adj 2.5×10^−6^), E2F targets (NES +1.86, p-adj 2.8×10^−5^), and nucleosome organization (NES +2.14, p-adj 9.1×10^−5^). In contrast, the eukaryotic translation initiation (NES −1.66, p-adj 0.016) and non-coding RNA-mediated heterochromatin formation (NES −1.92, p-adj 0.041) gene sets were downregulated (Fig. 1d,e). Rotation testing confirmed significant joint upregulation of genes from the mitotic cell cycle and E2F target sets, nucleosome organization, and heterochromatin formation (FDR 0.006-0.023). We then evaluated whether the significant gene sets shifted consistently in the expected direction in individual animals. For each set, we defined a module score as the mean z-scored expression, per animal, of its leading-edge (LE) genes, the genes that drive the enrichment signal. Module scores showed a consistent response per animal for representative pathways (Fig. 1f,g). Collectively, these findings demonstrate that ELS alters the transcriptional landscape of PND15 SPGs.

We then investigated whether ELS induced long-term transcriptional effects in spermatogonial cells. Adult SPGs (B2M⁻CD49f⁺CD90.2⁺) were FACS-enriched from adult control and ELS F1 males and profiled individually by RNA-seq. As in the PND15 libraries, deconvolution analysis estimated that SPGs accounted for over 98% of cells in every RNA-seq library, with minimal somatic signal and no compositional differences between groups (Supplementary Fig. 2j-l). PCA distinguished the two groups, indicating that ELS also affected the adult SPG transcriptome (Fig. 2a). We identified 23 differentially expressed genes (DEGs; FDR < 0.05; median absolute log2FC 0.88; Fig. 2b), with 12 upregulated and 11 downregulated (Fig. 2b,c). None of these genes overlapped with the DEGs at PND15, suggesting specific transcriptional effects. We next applied GSEA and found that the most significant gene sets in adult SPGs were largely downregulated (Fig. 2d,e). Gene sets involved in translation and RNA metabolism were among the most significantly downregulated, including mRNA splicing (NES −1.83, padj 2.4×10^−5^) and rRNA processing (NES −1.90, padj 1.7×10^−5^). Cell cycle mitotic (NES −1.52, padj 1.9×10^−4^), PRC2 methylation (NES −1.94, padj 2.2×10^−4^), and spermatogenesis (NES −2.01, padj 4.7×10^−7^) were also downregulated. Fewer sets were upregulated, including the RAC1 GTPase cycle (NES +1.72, padj 8.5×10^−4^) and GPCR signaling (NES +1.49, padj 0.0038) (Fig. 2d,e). Importantly, module scores showed a consistent response per animal for the representative pathways (Fig. 2f).

**Fig. 2.**
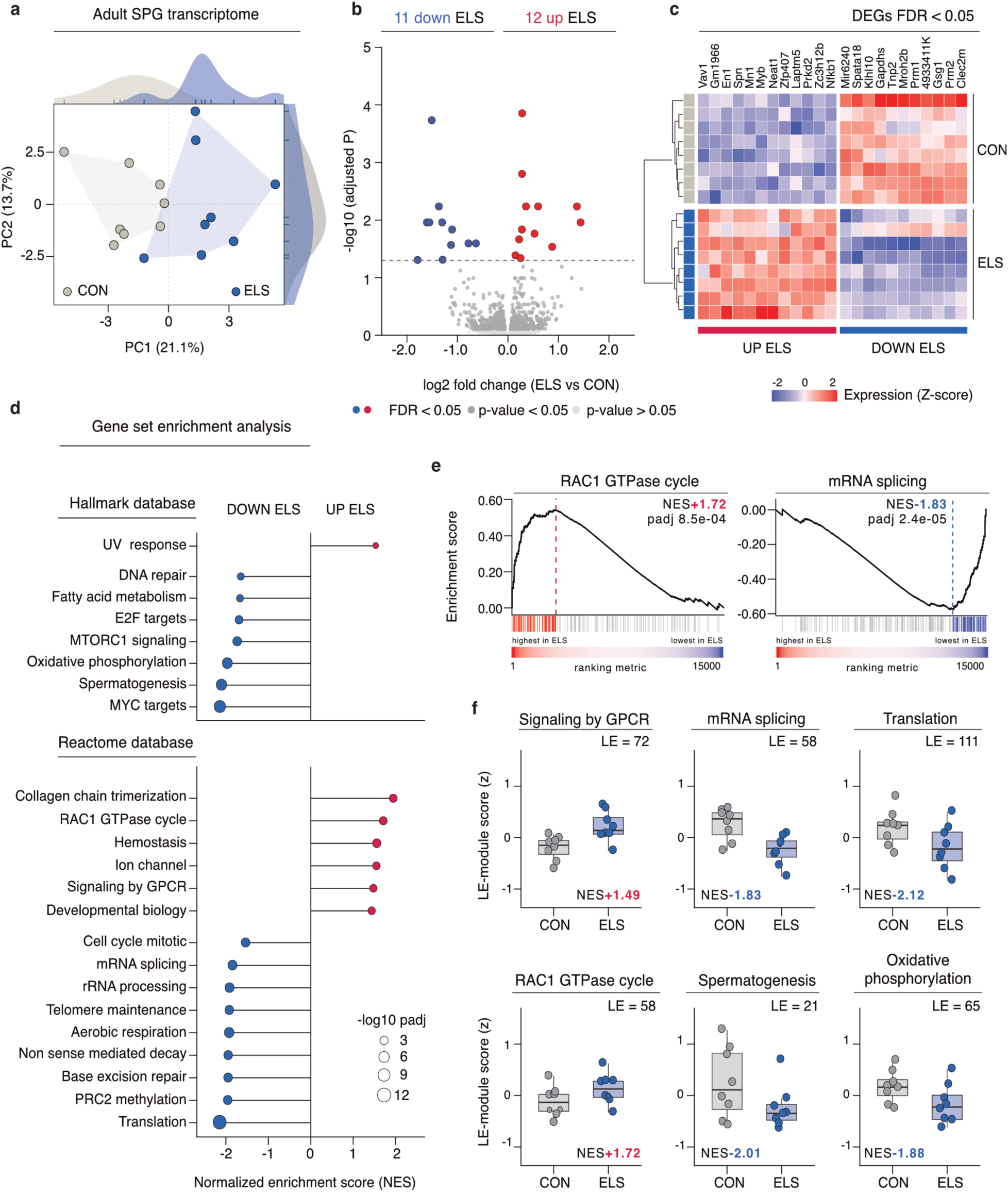
Early-life stress alters the spermatogonial transcriptome in adulthood. **a**, Principal component analysis of the SV-corrected transcriptome from spermatogonia (SPG) from control (CON) and early-life stress (ELS) adult F1 males. Axes show the variance explained by each component. *P* value by two-sided Wilcoxon rank-sum test on PC1 (CON *n* = 8; ELS *n* = 8 males; *P* = 0.0011). **b**, Volcano plot of differential gene expression between ELS and control SPGs. Color marks genes at FDR < 0.05 (red, higher in ELS; blue, lower), nominal *P* < 0.05 (mid grey), and *P* > 0.05 (light grey); counts above the panel give genes in each direction. *P*-values were obtained from the edgeR likelihood-ratio test adjusted for multiple testing (Benjamini–Hochberg) (CON *n* = 8; ELS *n* = 8 males). **c**, Heatmap of all 23 differentially expressed genes (FDR < 0.05), *z*-scored per gene and split by direction of change. Rows are individual males (CON *n* = 8; ELS *n* = 8), clustered (dendrogram at left) and annotated by group at right. **d**, Gene set enrichment analysis of the Hallmark and Reactome collections. Point position, normalized enrichment score (NES); point area, -log10 adjusted *P*; color, direction of change. All eight Hallmark sets at FDR < 0.05 are shown; Reactome sets are one representative per Reactome pathway-hierarchy group among the 124 sets at FDR < 0.05, ranked by adjusted *P*. **e**, Running enrichment score for RAC1 GTPase cycle (left) and mRNA splicing (right) across genes ranked by the signed likelihood-ratio statistic. Vertical lines mark gene set members; colored lines are the leading-edge (LE) genes for that set; the strip below shows the ranking metric from highest to lowest in ELS. **f**, LE module score per male for six enriched sets, computed on the log-CPM assay as the mean per-gene *z*-score of the leading-edge genes of each set. Box, interquartile range with median line; whiskers, 1.5× interquartile range; points, individual males (CON *n* = 8; ELS *n* = 8). Scores are not sign-adjusted, so sets lower in ELS sit below zero.

To further characterize ELS effects in the adult male germline, we profiled the whole adult testis using single-cell RNA-seq (scRNA-seq). Clustering of 54,745 cells identified thirteen cellular populations (Fig. 3a). We resolved eight cell types spanning the germline in the expected developmental sequence, from SSCs to differentiating spermatogonia (DiffSPGs), primary and secondary spermatocytes, early, mid, and late round spermatids, and elongating spermatids, consistent with previous single-cell testicular atlases^10,15,16,19^. The remaining five cell types comprised the somatic compartment, including Sertoli, myoid, endothelial, Leydig cells, and macrophages. *Zbtb16* and *Gfra1*, which mark undifferentiated spermatogonia^20,21^, were most highly expressed in the SSC cluster, confirming its identity (Fig. 3b). We did not observe any significant differences in cellular composition between groups (Wilcoxon rank-sum, all Benjamini-Hochberg-adjusted *P* = 1.0; Fig. 3c,d), indicating that ELS does not have a major impact on testicular cellular composition as measured by scRNA-seq.

**Fig. 3.**
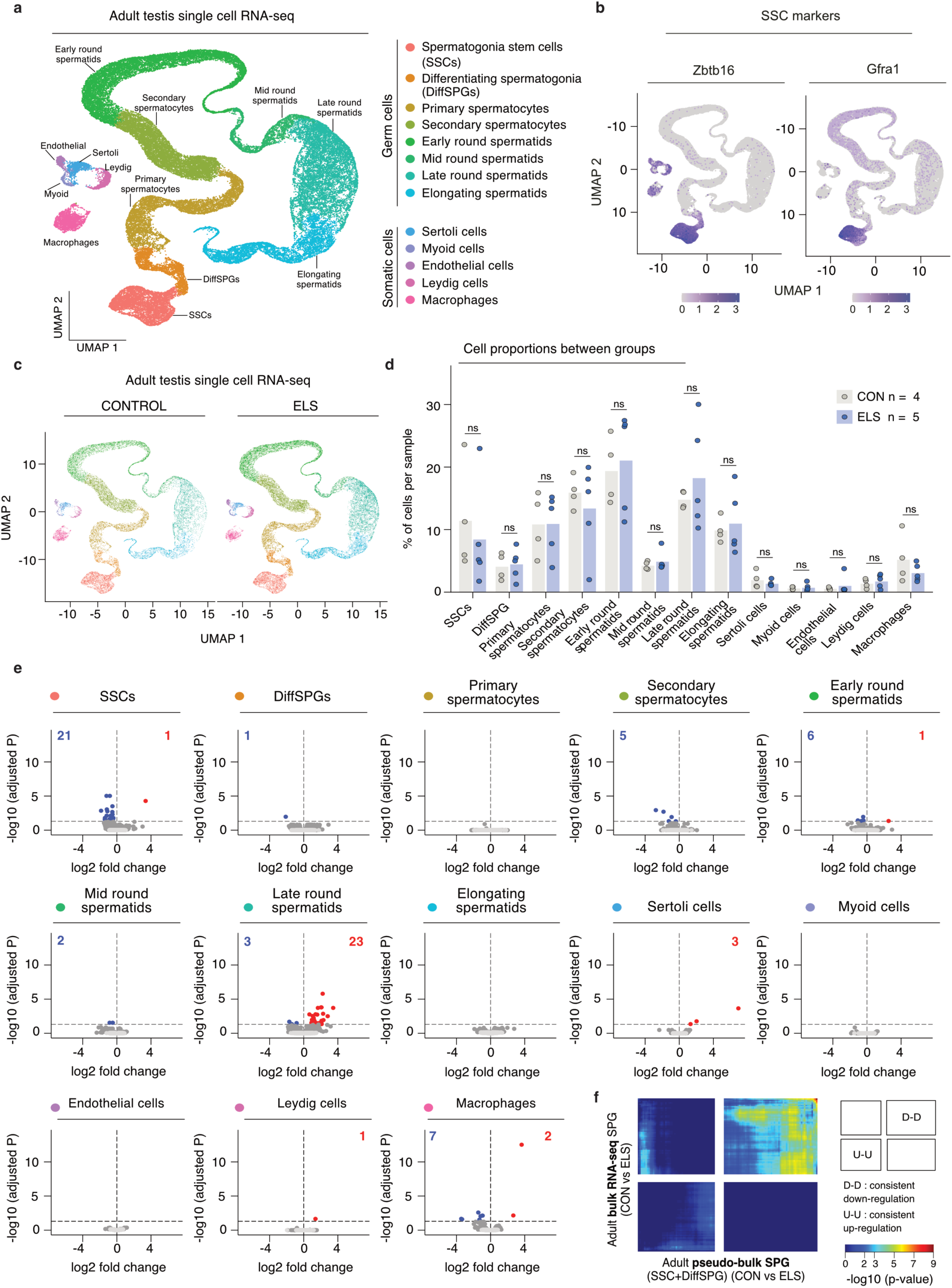
Early-life stress affects the adult testis transcriptome in a cell-type-restricted manner. **a**, UMAP of 54,745 single-cell transcriptomes from whole adult testis, coloured by annotated cell type; germ-cell and somatic populations are grouped in the key. Cells derive from control (CON) and early-life stress (ELS) F1 males of a cohort independent of the bulk RNA-seq datasets (CON *n* = 4; ELS *n* = 5 males; 3,478-7,006 cells per male). **b**, Expression of the spermatogonial stem cell (SSC) markers *Zbtb16* and *Gfra1* on the same embedding. **c**, UMAP split by condition, showing 22,529 control and 32,216 ELS cells pooled across males. **d**, Proportion of each annotated cell type per male. Each point is an individual male (CON *n* = 4; ELS *n* = 5); bars are means across males. *P* value by two-sided Wilcoxon rank-sum test adjusted for multiple testing (Benjamini–Hochberg) across the thirteen cell types; ns, adjusted *P* = 1.000 for all thirteen. **e**, Volcano plots of differential expression per cell type on a common scale. Color marks genes at FDR < 0.05 (red, higher in ELS; blue, lower); counts in the upper corners give genes in each direction. Dashed lines mark FDR = 0.05 and zero fold change. *P*-value by edgeR likelihood-ratio test on per-male pseudobulk profiles, adjusted for multiple testing (Benjamini–Hochberg) (CON *n* = 4; ELS *n* = 5 males). **f**, Rank–rank hypergeometric overlap between the pseudobulk spermatogonial ranking of this dataset (SSCs and DiffSPGs) merged; *x* axis; CON *n* = 4; ELS *n* = 5 males) and the adult bulk spermatogonial ranking of Fig. 2 (*y* axis; CON *n* = 8; ELS *n* = 8 males). Transcripts are ranked by −log10(*P*) × sign(log2 fold change) over the 12,716 transcripts detected in both datasets, in steps of 113. Color gives -log10 adjusted *P* of the overlap (Benjamini–Hochberg). D–D, transcripts lower in ELS in both datasets; U–U, higher in both. DiffSPG, differentiating spermatogonia; SSC, spermatogonial stem cell.

We then examined gene expression differences between control and ELS across multiple cell populations. We identified 76 genes differentially expressed at FDR < 0.05 across nine of thirteen cell types, with 45 genes downregulated and 31 upregulated (Fig. 3e). The SSC response was predominantly characterized by downregulated genes (21 downregulated genes vs 1 upregulated; median absolute log2FC 0.847). Notably, 16 of these 21 downregulated genes are involved in spermatogenesis and were highly expressed in round or elongating spermatids in this dataset. This aligns with the observed transcriptional downregulation of the spermatogenesis gene set in the Adult bulk RNA-seq (Fig. 2d). Among other cell types, late round spermatids exhibited the most transcriptional changes, with 23 genes upregulated and 3 downregulated. Among somatic cells, macrophages showed the most DEGs, with 7 downregulated and 2 upregulated. These findings suggest that ELS influences the testicular transcriptome in adult mice.

Finally, we examined whether the adult SPG bulk RNA-seq response could be detected in the single-cell data without setting a significance threshold. We combined the single-cell SSC and DiffSPG cell populations into a pseudo-bulk spermatogonial profile (SPG-sc) that mirrors the FACS-enriched fraction used for bulk RNA-seq, which includes both populations, with SSCs as the dominant group, as indicated by deconvolution analysis (Supplementary Fig. 2j-l). Next, we performed differential gene expression analysis on the SPG-sc pseudo-bulk datasets and ranked genes from most to least upregulated in ELS. Using rank-rank hypergeometric overlap, we tested whether the same DEGs appeared at similar positions in both the pseudo-bulk and bulk rankings. We found that the agreement was mainly limited to the shared downregulated genes (Fig. 3f). Elsewhere, the correlation between the datasets was no better than chance (Spearman’s ρ = 0.03). Overall, the data suggest that ELS causes both immediate and lasting changes in the SPG transcriptome.

### Early-life stress alters chromatin accessibility in spermatogonial cells

To characterize potential regulatory effects of ELS, we performed ATAC-seq on FACS-enriched SPGs from PND15 and adult control and ELS F1 males. Differential accessibility analysis at the individual-peak level identified just 28 genomic regions with differential chromatin accessibility in PND15 and none in adult (FDR < 0.05; data not shown). We therefore asked whether ELS instead produces coordinated, subtle shifts in chromatin accessibility across functionally related regions. To address this, we used MonaLisa^22^, which identifies transcription-factor motif enrichment along the continuum of accessibility change. We ranked consensus peaks by accessibility log2 fold change, divided them into ten equal-size bins, and tested motif enrichment per bin.

At PND15, peaks gaining accessibility in ELS were enriched for AP-1/bZIP family motifs (BACH2, FOS::JUNB, JDP2, NFE2, NFE2L1; max log2 enrichment +0.52 in the top bin, FDR < 0.001), whereas peaks losing accessibility were enriched for STAT motifs (STAT1/3/4/5A/5B; STAT1 bin-1 log2 enrichment +0.33, FDR < 0.001) and nuclear-receptor motifs (AR, NR3C1, NR3C2; bin-1 max log2 enrichment up to +0.59, FDR < 0.001) (Fig. 4a). To test whether these bin-level signals were representative of the full animal cohort, we examined accessibility at peaks with the identified motifs from the enriched bins per animal (Fig. 4b). At BACH2-motif peaks (n = 528, bin 10), accessibility was higher in ELS than in control animals in 527 of 528 peaks (Δmean +0.26 log2 CPM, Cohen’s *d* = 0.32; Kolmogorov-Smirnov *D* = 0.237, *P* = 2.81×10^−13^; Fig. 4b). At STAT1-motif peaks (n = 2,012, bins 1 and 3), accessibility was lower in ELS than in control animals in 1,977 of 2,012 peaks (Δmean −0.19 log2 CPM, Cohen’s d = −0.15; Kolmogorov-Smirnov D = 0.124, P = 8.28×10^−14^; Fig. 4b). Per-animal medians separated the groups in both sets, with every ELS animal lying above every control at BACH2 peaks and below every control at STAT1 peaks (Wilcoxon *P* = 0.0009 for both; Fig. 4b).

**Fig. 4.**
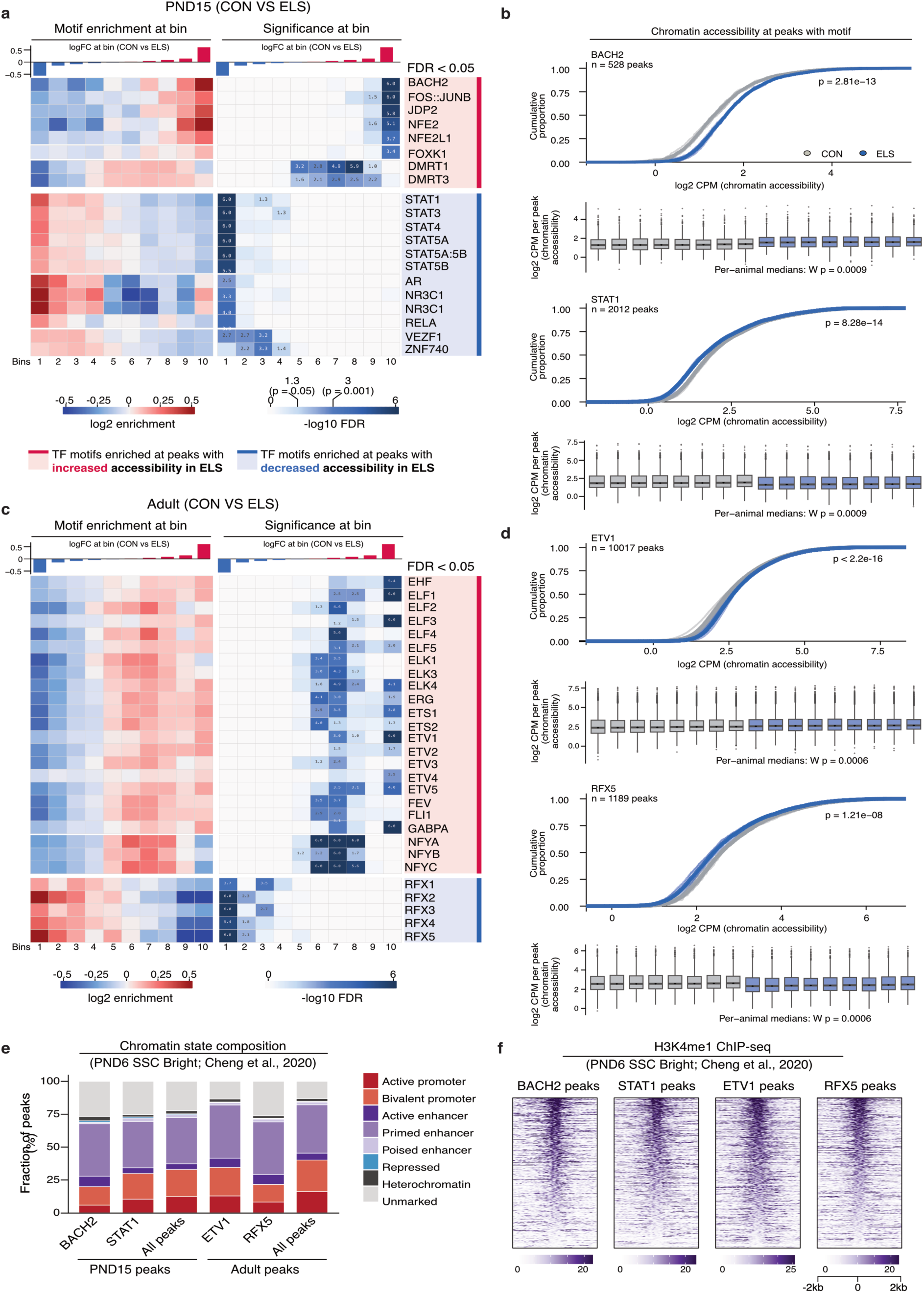
Early-life stress leaves age-specific transcription-factor motif signatures in spermatogonial chromatin accessibility. **a**, Transcription-factor motif enrichment across chromatin accessibility changes in spermatogonia (SPG) from control (CON) and early-life stress (ELS) F1 males at PND15. The 97,173 consensus peaks were ranked by log2 fold change (ELS versus control) and split into ten equal-occupancy bins (bin 1, strongest decrease; bin 10, strongest increase). Left, log2 motif enrichment per bin, color scale clipped at ±0.5; right, -log10 FDR, with scale and printed values capped at 6.0. Sidebars to the right: red, motifs enriched at peaks gaining accessibility in ELS; blue, motifs enriched at peaks losing accessibility. 20 of 41 robust motifs are shown; motifs enriched only in central bins that do not show changes in accessibility are removed for clarity (CON *n* = 8; ELS *n* = 8 males). **b**, Chromatin accessibility at motif-carrying peaks from the enriched bins in **a**. Top, cumulative distribution of log2 CPM per peak across 528 BACH2 peaks (bin 10) and 2,012 STAT1 peaks (bins 1 and 3); *P* value by two-sided Kolmogorov-Smirnov test on per-peak group means (BACH2 *P* = 2.81 × 10^−13^; STAT1 *P* = 8.28 × 10^−14^). Bottom, log2 CPM per peak with one box per male; box, interquartile range with median line; whiskers, 1.5× interquartile range; points beyond the whiskers, individual peaks. *P* value by two-sided Wilcoxon rank-sum test on per-male median accessibility (BACH2 *P* = 0.0009; STAT1 *P* = 0.0009; CON *n* = 8; ELS *n* = 8 males). **c**, As in **a** for adult SPGs, across 91,792 consensus peaks, with 28 of 57 robust motifs shown (CON *n* = 8; ELS *n* = 9 males). **d**, As in **b** for the adult signature, across 10,017 ETV1 peaks (bins 8-10) and 1,189 RFX5 peaks (bins 1-3). *P* value by two-sided Kolmogorov-Smirnov test (ETV1 *P* < 2.2 × 10^−16^; RFX5 *P* = 1.21 × 10^−8^) and by two-sided Wilcoxon rank-sum test on per-male median accessibility (*P* = 0.0006 for both; CON *n* = 8; ELS *n* = 9 males). **e**, Chromatin state composition of the four motif-defined peak sets and of their respective peak universes (All peaks), assigned from six histone marks in PND6 ID4-eGFP-Bright spermatogonia^23^. The reference epigenome is juvenile: for the adult sets it describes the developmental context of these regions, not their adult state. **f**, H3K4me1 ChIP-seq signal in the same reference dataset across ±2 kb of peak center in 10-bp bins, rows ordered by central signal, color capped at the 95th percentile per panel. ETV1 peaks were subsampled to 2,000. *n*, individual males; CPM, counts per million.

In adults, the TF motif signature was entirely distinct (Fig. 4c). None of the PND15 families (AP-1/bZIP, STAT, NR3C) were enriched. Instead, peaks gaining accessibility were enriched for ETS-family motifs, including ELK1, ETS1, ETV1, and FLI1, as well as for NFYA, NFYB, and NFYC. In contrast, peaks losing accessibility were enriched for RFX1-RFX5. At ETV1-motif peaks (n = 10,017, bins 8-10), accessibility was higher in ELS adults in 9,741 of 10,017 peaks (Δmean +0.18 log2 CPM, d = 0.16; D = 0.138, P < 2.2×10^−16^; Fig. 4d). At RFX5-motif peaks (n = 1,189, bins 1-3), accessibility was lower in ELS adults in 1,159 of 1,189 peaks (Δmean −0.19 log2 CPM, d = −0.18; D = 0.126, P = 1.21×10^−8^). Per-animal medians again completely separated the groups (Wilcoxon P = 0.0006 for both), supporting coordinated responses among the SPGs of individual males.

As an orthogonal precision control, we subjected the motif-accessibility associations to randomized lasso stability selection, which tests whether each motif is retained as an independent predictor of accessibility change in the presence of all others. At PND15, STAT1 was stably selected (selection probability >= 0.8), whereas the AP-1/bZIP signal fragmented across 45 nearly identical family variants (maximum selection probability 0.68), consistent with the high similarity among AP-1 TF motifs. At the adult timepoint, the ETS (ELK1 cluster), RFX (RFX5 cluster), and NFY (NFYA cluster) clusters were stably selected.

Finally, we annotated the four motif-defined peak sets against reference epigenomes derived from PND6 SPGs^23^. Most TF-associated peaks overlapped with regions annotated as enhancers, with primed enhancers accounting for 35-41% of peaks in each set (Fig. 4e). Consistent with the functional classification, we observed clear deposition of H3K4me1 at the four motif-defined peak sets in PND6 SPGs (Fig. 4f). Collectively, these data indicate that ELS redistributes accessibility at primed enhancers sharing specific TF motifs. As observed at the transcriptome level, the chromatin accessibility changes induced by ELS at PND15 and in adults are largely unique.

## DISCUSSION

Taken together, our data indicate that spermatogonial cells respond to early life stress at both the transcriptional and chromatin levels, and that a molecular signature of that exposure remains detectable in the adult spermatogonial pool. One day after the end of exposure, the transcriptional response centered on proliferation and chromatin programs, while regions that changed in chromatin accessibility were enriched for motifs of the AP-1/bZIP and STAT families of TFs. These families of TFs transduce systemic signals^24,25^, consistent with signaling input reaching SPGs during exposure. Interestingly, AP-1 and STAT TFs are involved in inflammatory memory in epithelial mammalian stem cells^26^, linking environmental exposures with lasting effects in chromatin accessibility. In adulthood, the spermatogonial transcriptional response was modest and directional, with single-cell and bulk RNA-seq detecting a prominent downregulated signal enriched for genes involved in spermatogenesis. At the chromatin level, regions that changed accessibility were enriched for the ETS, NFY, and RFX families of TFs. ETS TFs, such as ETV5, are important for SSC self-renewal^27^, and NFY TFs have been proposed to be relevant to SSCs and germ cell maturation^28^. The TF families enriched in adult SPGs were not among those enriched at PND15. SPGs undergo extensive transcriptional and chromatin remodeling between these two ages^29^, raising the possibility that the adult signature emerges through interaction with a different transcription-factor landscape during SPG maturation. Finally, whether the molecular state of SPGs after ELS influences the alterations reported in adult sperm remains to be determined.

## METHODS

### Experimental methods

#### Mouse husbandry and experimentation

Mice were housed in a temperature- and humidity-controlled facility under a reversed light–dark cycle. Food and water were provided ad libitum. All experiments were approved by the cantonal veterinary office, Zurich (license 34524 / ZH021/2022). For the ELS paradigm, C57BL/6 J primiparous females and males were mated at 2.5–3 months of age. Dams and litters were randomly assigned to the ELS or control group. Animals in the ELS group were exposed to 3 h of proximal, unpredictable separation combined with unpredictable maternal stress (MSUS) daily from PND1 to 14, as described previously. Animals in the control group were left undisturbed. Cages were changed once a week until weaning (PND21) for both ELS and control animals. At weaning, pups from the same experimental group and sex but different dams were assigned to social groups (4-5 mice per cage) to minimize litter effects.

#### Behavioral testing

Behavior was assessed using the elevated plus maze (EPM), as previously described^30^ with minor modifications. Briefly, mice approximately 2–2.5 months of age were singly housed for 1 h before testing and then transferred to the experimental room immediately before testing. The EPM consisted of two open and two closed arms (30 × 5 cm; closed-arm wall height, 15 cm) elevated 60 cm above the floor. Illumination was 18 ± 1 lux in the open arms and 9 ± 1 lux in the closed arms. Each mouse was placed on the central platform facing an open arm and allowed to explore the maze for 5 min. Time spent in the open and closed arms and total distance traveled were recorded using ViewPoint Behavior Technology. Latency to first enter an open arm was scored manually, and videos were subsequently reviewed to verify tracking accuracy. The experimenter was blinded to animal identity and treatment. Testing was conducted during the dark phase.

#### Fluorescence-activated cell sorting

For sorting PND15 SPG, testes were isolated, dealbulginated and digested in 2 ml Goni-MEM (DMEM (Gibco, 11965092) supplemented with 1x penicillin-streptavidin (Gibco, 15070063), 1x NEAA (Gibco, 11140050), 1x sodium pyruvate (Gibco, 11360070) and 1.68 µl ml^−1^ sodium lactate (Sigma-Aldrich, L4263)) with 0.5 mg ml^−1^ collagenase (Sigma-Aldrich, C7657) for 10-12 minutes at 32 °C, shaking at 1000 rpm on a thermomixer. The resulting seminiferous tubules were centrifuged at 200 rcf for 3 minutes at 4 °C and further digested in 1 ml 0.05% Trypsin–EDTA (Gibco, 25300054) supplemented with 10 µl 5 mg ml^−1^ DNase I (Sigma-Aldrich, DN25) for 10 minutes at 32 °C, shaking. The digestion was stopped by adding 200 µl FBS (Gibco, A5256701). An additional 20 µl of DNase I were added to the single-cell suspension and incubated for 3 minutes at 32 °C, shaking. Cells were centrifuged at 200 rcf for 3 minutes at 4 °C, resuspended in FACS buffer (PBS supplemented with 2% FBS) and centrifuged again. Next, cells were blocked with Fc block (1:50 anti-CD16/32, clone 93, eBioscience, 14-0161-81) in 100 µl FACS buffer for 20 minutes and labelled with anti-CD45 (1:400, clone 30-F11, eBioscience, 13-0451-82) and anti-CD51 (1:100 clone RMV-7, eBioscience, 13-0512-82) biotin conjugated antibodies in 200 µl FACS buffer supplemented with 0.01% NaN_3_ (Sigma-Aldrich, 08951) for 20 min. Cells were washed twice with 1 ml FACS buffer, and stained with anti-CD9^APC^ (1:200, clone KMC8, eBioscience, 17-0091-82), anti-c-Kit^PE-Cy7^ (1:1600, clone 2B8, eBioscience, 25-1171-81), streptavidin^V421^ (1:250, BD Horizon, 563259) in 100 µl FACS buffer supplemented with 0.01% NaN_3_ for 20 min. Before sorting, cells were again washed twice with 1 ml FACS buffer and DAPI (Invitrogen, 62248) was added at a concentration of 1 μg ml^−1^. CD9^+^CD45^−^CD51^−^cKIT^−^ cells were sorted either into DNA/RNA Shield (Zymo, R1200) for RNA-seq or PBS for ATAC-seq at 4 °C on a BD Aria III using a 85 µm nozzle. Cells sorted into DNA/RNA Shield were directly frozen on dry ice and stored at –80 °C until further processing, cells sorted into PBS were directly subjected to ATAC-seq library preparation. Lasers were operated at FSC 220 V, SSC 250 V, Pacific blue 400 V, PE-Cy7 400 V, APC 500 V.

For sorting adult SPG, testes were isolated, dealbulginated and digested in 20 ml Goni-MEM with 0.5 mg ml^−1^ collagenase for 10-12 minutes at 37 °C, shaking in a water bath. Seminiferous tubules were let to sediment for 5 minutes on ice and after removing the supernatant further digested in 5 ml 0.05% Trypsin–EDTA supplemented with 500 µl 5 mg ml^−1^ DNase I for 5-10 minutes at 37 °C. The digestion was stopped by adding 500 µl FBS. An additional 100 µl of DNase I were added to the single-cell suspension and incubated at room temperature for 2 min. Cells were passed through a pre-wetted 0.40 µm cell strainer (Falcon, 352340) into a new tube and another 5 ml Goni-MEM added to wash cell strainer before centrifugation at 600 rcf for 7 minutes at 4 °C. Cells were resuspended in 20 ml DPBS-S (DPBS (Gibco, 14190144), 1% FBS, 1x penicillin-streptavidin, 1x sodium pyruvate, 1 mg ml^−1^ glucose (Gibco, A2494001), 10 mM HEPES (Gibco, 15630106)), 5 ml each slowly pipetted onto 2ml of 30% percoll (percoll (MP, 219536980) in DPBS, 1% FBS, 1x penicillin-streptavidin) in 15 ml falcons, and centrifuged at 600 rcf for 8 minutes 4 °C with the brake turned off. Supernatant was removed and pellet resuspended in DPBS-S, combining again the 4 pellets per sample and centrifuged at 600 rcf for 5 minutes at 4 °C. Next, cells were incubated with primary antibody (1:100 biotin anti-CD49f, clone GoH3, BioLegend, 313603; 1:100 anti-CD90.2^PE^, clone 30-H12, BD Pharmingen, 553014; 1:100 anti-b2-microglobulin (B2M)^BV421^, clone S19.8, BD Pharmingen 744802) in 100 µl for 20 minutes and washed with 1 ml of DPBS-S before adding secondary antibody (1:200 Streptavidin Alexa Fluor 488, Invitrogen, S32354; 1:1000 LIVE/DEAD fixable far red dead cell stain, Life Technologies, L34973) and washed with 1 ml DPBS-S again. Stained cells were resuspended in FACS buffer and CD90.2^+^CD49f^+^B2m^−^ cells sorted either into DNA/RNA Shield for RNA-seq or PBS for ATAC-seq at 4 °C on a BD S6 using a 85 µm nozzle. Cells sorted into DNA/RNA Shield were directly frozen on dry ice and stored at –80 °C until further processing, cells sorted into PBS were directly subjected to ATAC-seq library preparation. Lasers were operated at FSC 225 V, SSC 250 V, Pacific blue 696 V, FITC 647 V, PE 573 V, APC 410 V.

For scRNA-seq SPG were isolated as for adult bulk samples described above, the gating was slightly adjusted. To additionally enrich for diploid and somatic cells the protocol was adjusted as follows: after the digestion with trypsin, 1/5 of the volume was removed, centrifuged at 400 rcf for 5 minutes at 4 °C, resuspended in 1 ml of 1 mg ml–1 hyaluronidase (Sigma-Aldrich, H6254) and incubated 5-10 minutes at 35 °C. After the incubation 6 ml of Goni-MEM were added and cell solution filtered through a 70 µm cell strainer (Falcon, 352350), before counting the cells and centrifuging at 400 rcf for 5 minutes at 4 °C. Cells were then resuspended in Goni-MEM at a concentration of 10^6^ cells per 100 µl, Hoechst 33342 (1:100, BD Pharmingen, 561908) and Mitotracker (1:1000, Invitrogen, M22426) were added and incubated in a waterbath at 35 °C for 30 minutes shaking. Cells were washed with 5 ml Goni-MEM, centrifuged at 400 rcf for 5 minutes at 4 °C and resuspended in FACS buffer before proceeding to cell sorting. Directly before sorting Sytox Orange (1:10000, Invitrogen, S34861) was added to distinguish dead cells. Per sample 10000 CD90.2^+^CD49f^+^B2m^−^(5000 strict gating, 5000 wide gating), 15000 diploid, sertoli enrichted FSC^high^ mitotracker^high^, and 35000 diploid cells were sorted into 1 tube. All cells for scRNA-seq were sorted into FACS buffer at 4 °C on a BD Aria III using a 100 µm nozzle. Lasers were operated at FSC 256V, SSC 263V, Pacific blue 696V, FITC 647V, PE 573V, and APC410V for SPGs and FSC 229V, SSC255V, Pacific blue 403V (day 1) or 493V (day 2), APC 218V, PE 483V for diploid cells.

#### RNA isolation

RNA from PND15 and adult sorted SPGs was isolated using the Quick-DNA/RNA Micropep Plus kit (Zymo, D7005) following the manufacturer’s recommendations for samples stabilized and stored in DNA/RNA shield, including in-column DNase treatment. RNA quality and quantity were assessed on a Bioanalyzer 2100 (Agilent) using the RNA 6000 Pico kit (Agilent, 5067-1513).

#### Bulk RNA sequencing

Libraries were prepared using the SMARTer Stranded Total RNA-Seq Kit v3 - Pico Input Mammalian (Takara, 120523) according to the manufacturer’s recommendations, with minor modifications. In brief, 5 ng of RNA was used as input per library and fragmented for 4 minutes. For rRNA depletion, the ZapR Mammalian rRNA Depletion kit (Takara, 634370) was used following the protocol described in the SMART-Seq Total RNA Library Prep with ZapR Depletion (with UMIs) sequencing kit (Takara, 634355). PCR1 was performed with 5 cycles, and PCR2 with 12. Libraries were pooled and sequenced on a DNBseq T7 in 100 bp paired-end.

#### Single-cell RNA sequencing

After FACS enrichment, cells were centrifuged at 600 rcf for 10 minutes at 4 °C. The supernatant was removed, and cells were resuspended in PBS. Libraries were prepared using the 10X Genomics 3’ Gene Expression kit v3.1 according to the manufacturer’s recommendations in 2 batches and sequenced on a NovaSeq X in paired-end mode at 150 bp.

#### ATAC sequencing

Per sample, 25000 sorted PND15 uSPG and 10000 adult uSPG were centrifuged at 500 rcf for 10 minutes at 4 °C. The supernatant was removed, and the pellets were resuspended in 50 µl of cold lysis buffer (10 mM Tris HCl pH 7.5, 10 mM NaCl (Invitrogen, AM9760G), 3 mM MgCl2 (Sigma-Aldrich, M1028), 0.1% NP-40 (Roche, 47479200), 0.1% Tween-20 (Sigma-Aldrich, P9416), 0.01% Digitonin (Sigma-Aldrich, 300410)). Lysates were incubated on ice for 3 minutes, then 1 ml of wash buffer (10 mM Tris HCl pH 7.5, 10 mM NaCl, 3 mM MgCl2, 0.1% Tween-20) was added. Samples were centrifuged at 500 rcf for 10 minutes at 4 °C. The supernatant was removed, and the pellets were gently resuspended in 50 µl of TD mix (1x of 2xTD buffer (20 mM Tris-HCl, pH 7.6, 10 mM MgCl2, 20% dimethylformamide (DMF, Sigma-Aldrich, 227056)), 0.1% Tween-20, 0.01% Digitonin, 2.5 µl 50 µl–1 Tn5 (Illumina Nextera XT, FC-131-1096), 10% nuclease-free water, PBS) and incubated for 30 minutes at 37 °C at 1000 rpm. DNA was isolated using a MinElute reaction cleanup kit (Qiagen, 28204), eluted in 10 µl of elution buffer, and stored at −20 °C until library preparation. For library generation, transposed DNA was amplified by PCR using NEBNext high-fidelity PCR master mix (NEB, M0541S) and 25 µM Illumina i5 and i7 primers (Illumina, FC-131-2001) with the following PCR protocol: 72 °C for 5 minutes, 98 °C for 30 seconds, then for PND15, 14 cycles, and for adult samples, 12 cycles of 98 °C for 10 seconds, 63 °C for 30 seconds, 72 °C for 1 minute, and hold at 4 °C. To remove primer dimers and untagmented fragments above 1000 bp, a double-sided cleanup with AMPure XP beads (Beckman Coulter, A63880) was performed. Libraries were mixed with 0.5x volume of room-temperature AMPure XP beads and incubated for 10 minutes, then separated using a magnetic rack and the supernatant transferred to a new tube. Supernatants were then mixed with 1.3x the original volume of AMPure XP beads, incubated for 10 minutes, and again separated using a magnetic rack. Beads were washed twice with fresh 80% ethanol, air-dried, resuspended in 20 µl nuclease-free water, and placed back on the magnetic rack. The supernatant containing the libraries was transferred to a new tube and stored at −20 °C until further use. Library quality and quantity were measured using Tapestation (Agilent) and Qubit (Thermo Fisher Scientific). Libraries were pooled and sequenced on a DNBseq T7 in 100 bp paired-end.

### Computational analysis

#### Bulk RNA-seq data processing and analyses

Read pairs were tagged with unique molecular identifiers (UMIs) using umi_tools extract (UMItools version 1.1.4) with parameters --bc-pattern NNNNNNNNCCCCCC and --extract-method=string. Raw sequencing reads were trimmed using trimGalore (version 0.6.7) with parameters -q 30, --length 30, and --stringency 2, and quality was assessed using FastQC (version 0.12.1) and FastQ Screen (version 0.15.2). For UMI-based deduplication, reads were aligned to the mm10 reference genome using STAR (version 2.7.10b), deduplicated using umi_tools dedup (UMItools version 1.1.4) with --paired and --buffer-whole-contig, and deduplicated reads were converted back to FASTQ format using samtools fastq (SAMtools version 1.17).

#### Read quantification

Gene-level counts were obtained using featureCounts (Subread package) in strand-specific mode (*-s 2*) against the mm10 RefSeq gene annotation.

#### Filtering and normalization

Genes were filtered for expression using *edgeR::filterByExpr* (edgeR version 3.36.0). Library-size normalization used the trimmed mean of M-values method implemented in *edgeR::calcNormFactors*.

#### Removal of unwanted variation

Unwanted technical variation was removed by surrogate variable analysis (sva version 3.42.0), implemented via *SEtools::svacor* (version 1.23.1).

#### Differential expression analysis

Differential expression was tested using edgeR (version 3.36.0). Gene-wise dispersions were estimated with *estimateDisp* and the treatment effect tested by likelihood-ratio test (*glmFit*, *glmLRT*) under the design ∼ Group + SVs. *P* values were adjusted using the Benjamini–Hochberg procedure, and genes with an adjusted *P* < 0.05 were considered differentially expressed.

#### Principal component analysis

Principal component analysis was performed on all tested genes, without selection for variable genes, on both the uncorrected and the surrogate-variable-corrected matrices. Group separation along each component was assessed by two-sided Wilcoxon rank-sum test.

#### Gene set enrichment analysis

Genes were ranked by the likelihood-ratio statistic signed by the direction of change. Rank-based enrichment was computed using fgsea (version 1.20.0) with 10,000 permutations and a fixed random seed, restricted to sets of 10 to 500 genes present in the ranking. Three collections were retrieved with msigdbr (version 26.1.0) for *Mus musculus*: the mouse Hallmark collection, the Reactome canonical pathway subset, and Gene Ontology biological process.. Every set was additionally tested using two correlation-aware methods from limma (version 3.50.3): *camera*, with the inter-gene correlation estimated from the data and applied across each complete collection, and *fry*, a rotation test applied to *voom*-transformed counts.

#### Reduction of redundancy among enriched sets

Enriched sets were collapsed by two criteria. Sets were clustered on the pairwise containment of their leading-edge gene lists, defined as the size of the intersection divided by the size of the smaller list, using complete linkage at a threshold of 0.8. Significant Reactome pathways were separately grouped by ancestry in the Reactome hierarchy, with pathways assigned to the same group when one is an ancestor of the other, and each group labelled by its most significant member. Sets shown in Figure 1 were selected from the union of the two groupings after inspection of leading-edge composition.

#### Module scores

Per-animal module scores were computed as the mean gene-wise *z*-score of the leading-edge genes of a set on the surrogate-variable-corrected matrix. Scores were not adjusted by the sign of the normalised enrichment score. No statistical test was applied, as leading-edge genes are selected on the treatment contrast.

#### Data visualization

Volcano plots show log2 fold change against negative log10 adjusted *P* value, with genes coloured by adjusted *P* < 0.05, nominal *P* < 0.05, or neither. Heatmaps show gene-wise *z*-scores of the surrogate-variable-corrected matrix. Hierarchical clustering used Ward’s D2 linkage on Euclidean distances, and *z*-scores were clipped at ±2 for color after clustering. In the heatmap of differentially expressed genes, samples were clustered and genes ordered by functional category; in the leading-edge heatmap, samples were clustered and genes ordered by rank in the ranked gene list. Module scores are shown as box plots, with the box spanning the interquartile range, a line at the median, whiskers extending to 1.5 times the interquartile range, and individual animals plotted as points. Enrichment plots show the running enrichment score across the ranked gene list, with set members marked beneath and the ranking metric shown as a color gradient. In the gene set summary plot, point position gives the normalized enrichment score and point size the negative log10 adjusted *P* value.

#### Testis single-cell RNA-seq data processing and analyses

Single-cell RNA-seq data were processed with Cell Ranger count (10x Genomics Cell Ranger version 8.0.0) using the mouse mm10 reference transcriptome and --create-bam=true. Gene-barcode matrices were imported into Seurat (version 5.4.0). Ambient RNA was corrected with SoupX (version 1.6.2), and doublets were identified with scDblFinder (version 1.8.0) and removed. Cells were retained if they had 500–57,000 UMIs, 250–9,350 detected genes, and <10% mitochondrial reads. Genes detected in fewer than 10 cells were excluded. Data were normalized and integrated using the SCTransform workflow in Seurat, regressing out mitochondrial percentage and integrating batches with 2,000 variable features. Dimensionality reduction was performed with PCA, and UMAP was generated from the first 15 principal components. Clustering was performed with Seurat FindNeighbors and FindClusters. Cell types were assigned based on canonical marker expression. For pseudobulk differential expression analysis, raw RNA counts were aggregated by cell type, group, and sample using Seurat AggregateExpression. Differential expression was performed with edgeR (version 3.36.0) with experimental group as the variable of interest and batch as a covariate. Genes with FDR-adjusted P < 0.05 were considered differentially expressed. Additional analyses and visualization used Matrix (version 1.6-4), ggplot2 (version 3.5.2), dplyr (version 1.2.1), readr (version 2.2.0), SEtools (version 1.23.1), and pheatmap (version 1.0.13).

#### Deconvolution Analysis

Hermann et al., 2018: Raw data were obtained from adult sorted ID4-EGFP+SPGs. Using the cluster IDs provided in the supplemental datasets, CIBERSORTx analysis was performed to estimate the proportion of labeled cell types in each RNA-seq library using the default settings (G.min=300, G.max=500, q=0.1). For deconvolution analyses, we used the aggregated adult datasets as references. Tan et al., 2020: Raw data from E18, P2, and P7 were obtained. We applied the reported filtering and normalization parameters, followed by UMAP analysis (Becht et al., 2019) using the top 10 principal components (PCs). For clustering, the Seurat package was employed (Butler et al., 2018, dims = 1:10 in FindNeighbors, resolution = 0.025 in FindClusters) and resultant cell counts in each cluster corresponded to those reported in Tan et al., 2020. Clusters identified as germ, Sertoli, stroma, Leydig and peritubular myoid (PTM) cells were validated using specific marker genes as per Tan et al., 2020; Green et al., 2018; Hermann et al., 2018. Next, CIBERSORTx was utilized to estimate the proportion of these labeled cell types in each RNA-seq library using the default settings (G.min=300, G.max=500, q=0.1). For the deconvolution analyses, two replicate libraries from PND7 were used as references, and their averaged results were reported. Green et al., 2018: Raw data from adult were obtained. We applied the reported filtering and normalization parameters, followed by UMAP analysis (Becht et al., 2019) using the top 10 PCs. For clustering, the Seurat package (Butler et al., 2018, dims = 1:20 in FindNeighbors, resolution = 0.011 in FindClusters). Clusters identified as germ, Sertoli, stroma, Leydig, and peritubular myoid (PTM) cells were validated using specific marker genes as per Tan et al., 2020; Green et al., 2018; Hermann et al., 2018. Next, CIBERSORTx was utilized to estimate the proportion of these labeled cell types in each RNA-seq library using the default settings (G.min=300, G.max=500, q=0.1). For the final analysis, we used eight seminiferous tubule (ST datasets) and the averages of each group were reported.

#### Spermatogonia ATAC-seq data processing

Raw sequencing reads were trimmed using trimGalore (version 0.6.7) with parameters -q 30, --length 30, and --stringency 2, and quality was assessed using FastQC (version 0.12.1) and FastQ Screen (version 0.15.2). Reads were aligned to the mm10 reference genome using Bowtie2 (version 2.5.1) with parameters -X 2000, --end-to-end, and --very-sensitive. Duplicate reads, mitochondrial reads, and reads overlapping ENCODE blacklist regions were removed. Reads were shifted to account for Tn5 integration bias using alignmentSieve (deepTools version 3.5.4) with --ATACshift option, and nucleosome-free (NF) fragments were selected by removing fragments longer than 147 bp.

#### Peak calling and consensus peak sets

Peaks were called per sample on shift-corrected alignments using MACS2 (version 2.2.7.1) *callpeak* with parameters -f BAMPE -g mm --nomodel --keep-dup all -q 0.05. For each age, a consensus set was built as the union of per-sample peaks, retaining peaks detected in at least two samples, and each consensus peak was re-sized as a 501-bp window centered on the peak summit (summit +/− 250 bp). This yielded 97,173 windows at PND15 (n = 8 control vs 8 ELS animals) and 91,792 windows in adults (n = 8 vs 9).

#### Quantification, normalisation and removal of unwanted variation

NF fragments were counted per window, and library sizes were normalised by the trimmed mean of M-values method (*edgeR::calcNormFactors*; edgeR version 3.36.0). Unwanted variation was estimated with RUVr (RUVSeq version 1.28.0, k = 2): factors of unwanted variation (W) were computed from the deviance residuals of a first-pass edgeR fit under a group-only design, using all windows as control features. A batch-corrected log2 counts-per-million matrix, used for per-peak visualisation and the peak-set analyses below, was obtained with *limma::removeBatchEffect* (limma version 3.50.3) with the two W factors as covariates and the group design retained.

#### Differential accessibility

Differential accessibility between ELS and control animals was tested per window using edgeR under the design ∼ Group + W1 + W2. Dispersions were estimated with *estimateDisp*, and the group effect was tested with the quasi-likelihood framework (*glmQLFit*, *glmQLFTest*) on the group coefficient. P values were adjusted using the Benjamini-Hochberg procedure. Per-window log2 fold changes from this model were carried forward to motif analysis.

#### Binned transcription-factor motif enrichment

Motif enrichment along the continuum of accessibility change was computed with monaLisa (version 1.0.0). Windows were ranked by differential-accessibility log2 fold change and divided into ten equal-occupancy bins (bin 1, strongest decrease in ELS; bin 10, strongest increase). Per-bin enrichment of JASPAR2020 vertebrate motifs (JASPAR2020 version 0.99.10, retrieved with *TFBSTools::getMatrixSet*, TFBSTools version 1.32.0) was tested with *calcBinnedMotifEnrR* using Fisher’s exact test against all remaining bins as background, and P values were adjusted using the Benjamini-Hochberg procedure. To guard against parameter-dependent results, the analysis was repeated across a systematic grid of configurations varying the peak window type (fixed- and variable-width 501 bp), differential model (edgeR; additionally limma in adults), motif set, minimum absolute log2 fold-change filter, and bin number (10, 15 or 20), for a total of 55 configurations at PND15 and 110 in adults. Motifs were classified as robust when enriched at adjusted P < 0.05 in at least 80% of configurations within each peak-window by motif-set stratum, and were required to be robust in both window types (41 motifs at PND15; 57 in adults). Figure 4a,c displays robust motifs with directionally consistent enrichment in the extreme bins; motifs whose enrichment was confined to regions without change in accessibility (central bins) were not used for further analysis. Heatmap cell values are shown capped at |log2 enrichment| = 0.5.

#### Motif-carrying peak sets and accessibility analyses

Motif hits within the 501-bp windows were identified with *monaLisa::findMotifHits* (min.score = 10). For each transcription factor shown in Fig. 4b,d, the peak set comprised motif-carrying windows from that factor’s enriched bins (BACH2, bin 10; STAT1, bins 1 and 3; ETV1, bins 8-10; RFX5, bins 1-3). Cumulative distributions show batch-corrected log2 counts per million per peak, compared between groups by two-sample two-sided Kolmogorov-Smirnov test on per-peak group means. Mean differences are reported on per-peak group means, and Cohen’s d was computed as the pooled two-sample effect size over all per-peak, per-animal values, and is likewise descriptive. Group inference was based on per-animal statistics: for each animal, the median batch-corrected log2 counts per million across the set’s peaks was computed, and groups were compared by two-sided Wilcoxon rank-sum test with the normal approximation. Linear mixed models (*lmerTest::lmer*, version 3.2.1; log2 CPM ∼ group + (1 | animal), Satterthwaite degrees of freedom) were fitted as a complementary peak-level test; at PND15 these models were singular and are reported as such, and inference at this age rests on the per-animal tests. Box plots follow the conventions described above (box, interquartile range; line, median; whiskers, 1.5 times the interquartile range).

#### Chromatin-state annotation of peak sets

Reference epigenomic data of PND6 ID4-eGFP-Bright spermatogonia were obtained from Cheng et al. 2020^23^ (GEO GSE131656, ChIP-seq for H3K4me1, H3K4me2, H3K4me3, H3K27ac, H3K27me3 and H3K9me3; GSE131655, ATAC-seq; GSE131654, MeDIP-seq), using the processed coverage tracks and MACS2 peak calls provided. Each consensus window was scored for binary overlap (>= 1 bp, *GenomicRanges::findOverlaps*, GenomicRanges version 1.46.1) with each histone-mark peak set and assigned a chromatin state by sequential rules following Cheng et al.: repressed (H3K27me3 only), primed enhancer (H3K4me1 without H3K27ac or H3K27me3), poised enhancer (H3K4me1 with H3K27me3), active enhancer (H3K4me1 with H3K27ac, without H3K4me3), bivalent promoter (H3K4me3 with H3K27me3), active promoter (H3K4me3 with H3K27ac), heterochromatin (H3K9me3, assigned last), and unmarked otherwise. As the reference epigenome derives from juvenile spermatogonia, this annotation describes the developmental chromatin context of the PND15 and adult peak at PND6 rather than their actual state.

#### Reference ChIP-seq signal heatmaps

H3K4me1 coverage (PND6 Bright) was extracted in 10-bp bins across +/− 2 kb around each window centre (*rtracklayer*, version 1.54.0). Heatmap rows were ordered by mean signal in the central +/− 250 bp, and the colour scale was capped at the 95th percentile per panel. The ETV1 set was subsampled to 2,000 windows with a fixed random seed for display.

#### Statistical analysis

All statistical tests were two-sided, and P values were adjusted for multiple testing using the Benjamini-Hochberg procedure where indicated. Analyses were performed in R (version 4.1.2) with Bioconductor (release 3.14).

## AUTHOR CONTRIBUTIONS

RGA-M: Conceptualization, Investigation, Data curation, Software, Formal analysis, Visualization, Methodology, Supervision, Writing.

TS: Investigation, Formal analysis, Methodology, Writing. KU: Data curation, Software, Formal analysis, Methodology. IL-C: Conceptualization, Investigation.

IMM: Conceptualization, Resources, Supervision, Funding acquisition, Investigation, Visualization, Methodology, Writing, Project administration.

## ACKNOWLEDGMENTS

We thank Anna Chamot, Julio Eduardo Cáceres Pajuelo and Francesca Manuella for taking care of animal breeding and phenotyping. Leonardo Zingler Herrero and Leonard Steg for their support during FACS. The Mansuy lab is funded by the University Zurich, ETH Zurich, the Swiss National Science Foundation (grant number 31003A_175742/1), the National Centre of Competence in Research RNA&Disease funded by the Swiss National Science Foundation (grant numbers 182880/Phase 2 and 205601/Phase 3), ETH grants (ETH-10 15-2 and ETH-17 13-2), the Hochschulmedizin Flagship Project “STRESS,” the European Union Horizon 2020 Research Innovation Program EarlyCause (grant number 848158), the Horizon Europe program Staying Healthy 2021 under grant agreement number 101057529 (FAMILY) and grant agreement number 101057390 (HappyMums) funded by the Swiss State Secretariat for Education, Research and Innovation, ERA-NET NEURON Project EMPATHY (funded by the European Union’s Horizon 2020 research and innovation program under grant agreement number 964215), FreeNovation grant from Novartis Forschungsstiftung, and the Escher Family Fund. R.G.A.-M. received an ETH Postdoctoral Fellow grant (grant number 20-1 FEL-28).

## DATA AND MATERIALS AVAILABILITY

All datasets in this study will be made publicly available upon acceptance of the manuscript. All genomic raw and processed datasets will be deposited in the National Center for Biotechnology Information Gene Expression Omnibus (NCBI GEO).

## REFERENCES

1. Madigan, S. et al. Adverse childhood experiences: a meta-analysis of prevalence and moderators among half a million adults in 206 studies. World Psychiatry 22, 463–471 (2023).

2. Hughes, K. et al. The effect of multiple adverse childhood experiences on health: a systematic review and meta-analysis. Lancet Public Health 2, e356–e366 (2017).

3. Teissier, A. et al. Early-life stress impairs postnatal oligodendrogenesis and adult emotional behaviour through activity-dependent mechanisms. Mol. Psychiatry 25, 1159–1174 (2020).

4. Yam, K. Y. et al. Exposure to chronic early-life stress lastingly alters the adipose tissue, the leptin system and changes the vulnerability to western-style diet later in life in mice. Psychoneuroendocrinology 77, 186–195 (2017).

5. Franklin, T. B. et al. Epigenetic transmission of the impact of early stress across generations. Biol. Psychiatry 68, 408–415 (2010).

6. Gapp, K. et al. Implication of sperm RNAs in transgenerational inheritance of the effects of early trauma in mice. Nat. Neurosci. 17, 667–669 (2014).

7. Sharma, U. et al. Biogenesis and function of tRNA fragments during sperm maturation and fertilization in mammals. Science 351, 391–396 (2016).

8. Argaw-Denboba, A. et al. Paternal microbiome perturbations impact offspring fitness. Nature 629, 652–659 (2024).

9. Bellvé, A. R. et al. Spermatogenic cells of the prepuberal mouse. Isolation and morphological characterization. J. Cell Biol. 74, 68–85 (1977).

10. Ernst, C., Eling, N., Martinez-Jimenez, C. P., Marioni, J. C. & Odom, D. T. Staged developmental mapping and X chromosome transcriptional dynamics during mouse spermatogenesis. Nat. Commun. 10, 1251 (2019).

11. Drumond, A. L., Meistrich, M. L. & Chiarini-Garcia, H. Spermatogonial morphology and kinetics during testis development in mice: a high-resolution light microscopy approach. Reproduction 142, 145–155 (2011).

12. de Rooij, D. G. The nature and dynamics of spermatogonial stem cells. Development 144, 3022–3030 (2017).

13. Kubota, H. & Brinster, R. L. Spermatogonial stem cells. Biol. Reprod. 99, 52–74 (2018).

14. McCarrey, J. R. Toward a more precise and informative nomenclature describing fetal and neonatal male germ cells in rodents. Biol. Reprod. 89, 47 (2013).

15. Green, C. D. et al. A Comprehensive Roadmap of Murine Spermatogenesis Defined by Single-Cell RNA-Seq. Dev. Cell 46, 651–667.e10 (2018).

16. Hermann, B. P. et al. The Mammalian Spermatogenesis Single-Cell Transcriptome, from Spermatogonial Stem Cells to Spermatids. Cell Rep. 25, 1650–1667.e8 (2018).

17. Oatley, J. M. & Brinster, R. L. Regulation of spermatogonial stem cell self-renewal in mammals. Annu. Rev. Cell Dev. Biol. 24, 263–286 (2008).

18. van Steenwyk, G., Roszkowski, M., Manuella, F., Franklin, T. B. & Mansuy, I. M. Transgenerational inheritance of behavioral and metabolic effects of paternal exposure to traumatic stress in early postnatal life: evidence in the 4th generation. Environ. Epigenetics 4, dvy023 (2018).

19. Tan, K., Song, H.-W. & Wilkinson, M. F. Single-cell RNAseq analysis of testicular germ and somatic cell development during the perinatal period. Development 147, dev183251 (2020).

20. Costoya, J. A. et al. Essential role of Plzf in maintenance of spermatogonial stem cells. Nat. Genet. 36, 653–659 (2004).

21. He, Z., Jiang, J., Hofmann, M.-C. & Dym, M. Gfra1 silencing in mouse spermatogonial stem cells results in their differentiation via the inactivation of RET tyrosine kinase. Biol. Reprod. 77, 723–733 (2007).

22. Machlab, D. et al. monaLisa: an R/Bioconductor package for identifying regulatory motifs. Bioinformatics 38, 2624–2625 (2022).

23. Cheng, K. et al. Unique Epigenetic Programming Distinguishes Regenerative Spermatogonial Stem Cells in the Developing Mouse Testis. iScience 23, 101596 (2020).

24. Vierbuchen, T. et al. AP-1 Transcription Factors and the BAF Complex Mediate Signal-Dependent Enhancer Selection. Mol. Cell 68, 1067–1082.e12 (2017).

25. Bejjani, F., Evanno, E., Zibara, K., Piechaczyk, M. & Jariel-Encontre, I. The AP-1 transcriptional complex: Local switch or remote command? Biochim. Biophys. Acta BBA - Rev. Cancer 1872, 11–23 (2019).

26. Larsen, S. B. et al. Establishment, maintenance, and recall of inflammatory memory. Cell Stem Cell 28, 1758–1774.e8 (2021).

27. Ishii, K., Kanatsu-Shinohara, M., Toyokuni, S. & Shinohara, T. FGF2 mediates mouse spermatogonial stem cell self-renewal via upregulation of Etv5 and Bcl6b through MAP2K1 activation. Development 139, 1734–1743 (2012).

28. Dolfini, D., Imbriano, C. & Mantovani, R. The role(s) of NF-Y in development and differentiation. Cell Death Differ. 32, 195–206 (2025).

29. Lazar-Contes, I. et al. Dynamics of transcriptional programs and chromatin accessibility in mouse spermatogonial cells from early postnatal to adult life. eLife 12, RP91528 (2025).

30. Boscardin, C., Manuella, F. & Mansuy, I. M. Paternal transmission of behavioural and metabolic traits induced by postnatal stress to the 5th generation in mice. Environ. Epigenetics 8, dvac024 (2022).

