## Supplemental Figures for "Early life stress affects the transcription and chromatin accessibility of spermatogonial cells"

**Supplementary Fig. 1**

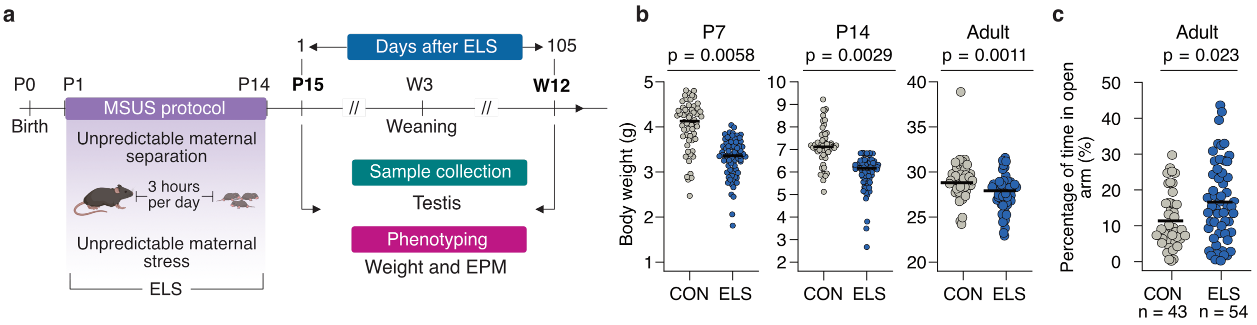


**Supplementary Fig. 1. Early-life stress reduces body weight and increases open-arm exploration in F1 males. a**, Experimental design. Early-life stress (ELS) was induced in C57BL/6J mice by unpredictable maternal separation combined with unpredictable maternal stress (MSUS): F1 pups were separated from the F0 dam for 3 h daily at unpredictable times during the dark phase from PND1 to PND14, and during each separation the dam was subjected, randomly and unpredictably, to either a 5-min forced swim in cold water (18 °C) or 20-min restraint in a tube. Dams and pups were housed in separate cages within visual and olfactory contact. Control (CON) dams and litters were left undisturbed apart from weekly cage changes. Litters were undisturbed from PND15 and weaned at PND21 (W3). Testes were collected from F1 males at PND15, one day after the end of the protocol, and at week 12 (W12), 105 days later. All phenotypes shown here were measured in F1 males from the same cohort as the males analyzed in Figs. 1, 2, and 4; the single-cell dataset in Fig. 3 comes from a separate cohort. Mouse cartoon created with BioRender. **b**, Body weight of F1 males at PND7, PND14 and in adulthood. Each point is an individual male (PND7 and PND14, $n$ = 56 control and 68 ELS; adult, $n$ = 44 control and 53 ELS); bars are medians, and the $y$-axis is independent in each panel. $P$ values at PND7 and PND14 were computed on litter means ($N$ = 14 control and 14 ELS litters). $P$ value by two-sided Wilcoxon rank-sum test, Holm-adjusted across the time points assessed (PND7 $P$ = 0.0058; PND14 $P$ = 0.0029; adult $P$ = 0.0011). **c**, Percentage of the total test duration spent in the open arms of the elevated plus maze, in F1 males at week 8. Each point is an individual male ($n$ = 43 control and 54 ELS); bars are means. $P$ value by two-sided Wilcoxon rank-sum test ($P$ = 0.023, unadjusted). $n$, individual males.

**Supplementary Fig. 2**

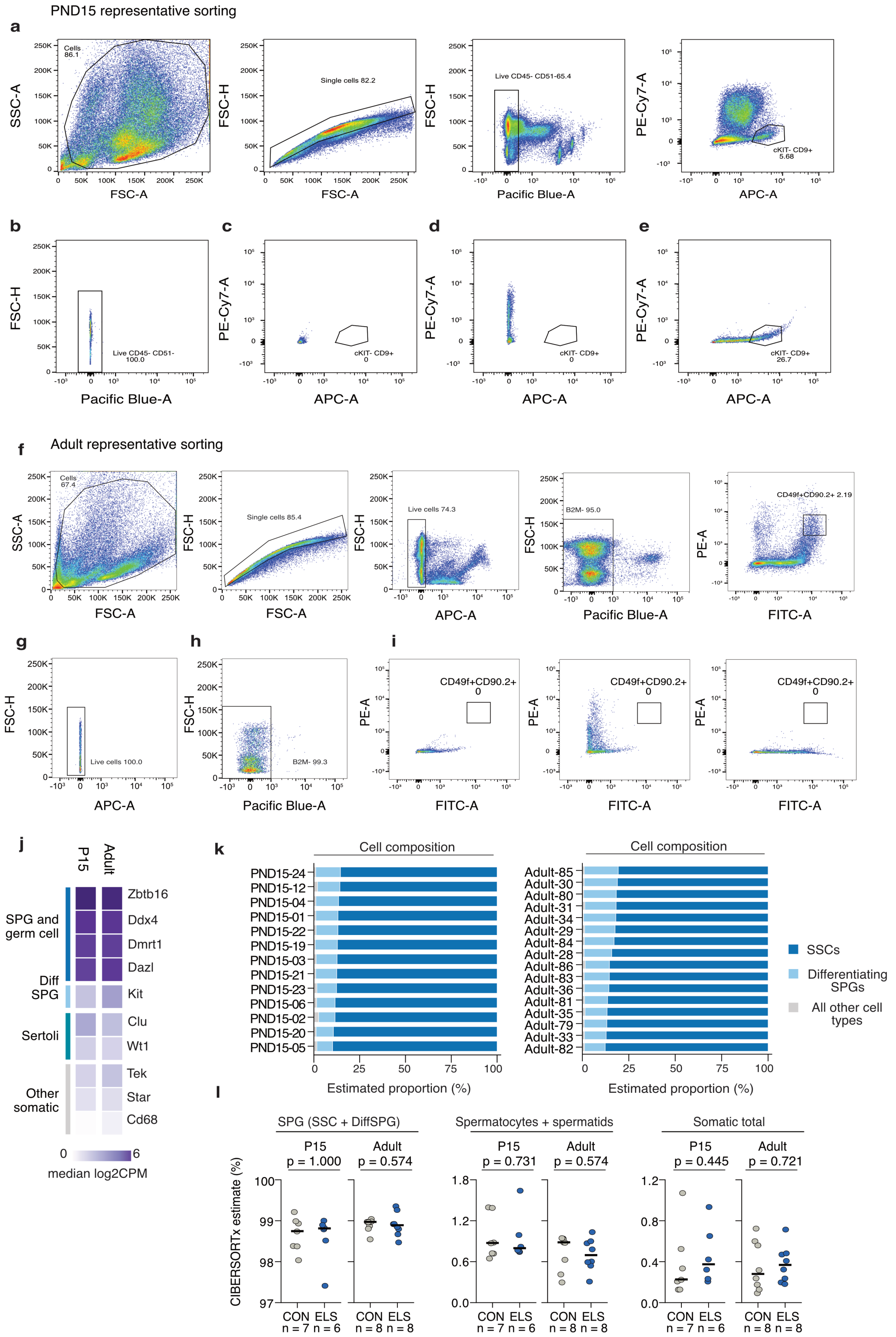

**Supplementary Fig. 2. Sorting strategy and transcriptomic validation of spermatogonial identity. a**, Representative gating for one PND15 F1 male: cells (FSC-A/SSC-A), single cells (FSC-A/FSC-H), live CD45−CD51− cells (Pacific Blue-A/FSC-H) and the sorted CD9+cKIT− spermatogonia (SPG; APC-A/PE-Cy7-A). Percentages give the frequency of the parent population, here and in **b**–**i**. **b**–**e**, Control stains defining the PND15 gate boundaries: **b**, unstained control for the live CD45−CD51− gate; **c**, unstained and **d**, single-stain controls for the CD9/cKIT gate; **e**, full-stain control. **f**, Representative gating for one adult F1 male: cells (FSC-A/SSC-A), single cells (FSC-A/FSC-H), live cells (APC-A/FSC-H), B2M− cells (Pacific Blue-A/FSC-H) and the sorted CD49f+CD90.2+ SPG (FITC-A/PE-A). **g**–**i**, Control stains defining the adult gate boundaries: **g**, unstained control for the live-cell gate; **h**, unstained control for the B2M− gate; **i**, controls for the CD49f/CD90.2 gate. **j**, Median log2 CPM across F1 males for lineage markers at PND15 ($n$ = 13) and in adulthood ($n$ = 16). Spermatogonial and pan-germ-cell markers (*Zbtb16*, *Ddx4*, *Dmrt1*, *Dazl*) are among the most abundant transcripts at both ages (64–126 CPM), whereas Sertoli and other somatic markers (*Clu*, *Wt1*, *Tek*, *Star*, *Cd68*) are an order of magnitude or more lower (1.7–11 CPM). The differentiation marker *Kit* is low at both ages and higher in adults (6.0 versus 13.4 CPM), consistent with sorting for the undifferentiated compartment. **k**, Estimated cellular composition of each library by CIBERSORTx deconvolution against a testis single-cell reference of thirteen cell types, one bar per male, ordered by estimated spermatogonial stem cell (SSC) proportion. Spermatogonia (SSC plus differentiating spermatogonia, DiffSPG) account for 97.4-99.2% of PND15 and 98.5–99.4% of adult libraries, leaving 0.8-2.6% and 0.6-1.5% for all other cell types. **l**, Estimated proportions per male in control (CON) and early-life stress (ELS) males, for three mutually exclusive compartments together covering each library: spermatogonia, spermatocytes plus spermatids, and total somatic cells (Sertoli, peritubular myoid, endothelial and Leydig). Each point is an individual male; bars are medians. $P$ value by two-sided Wilcoxon rank-sum test, unadjusted (PND15, CON $n$ = 7; ELS $n$ = 6 males; adult, CON $n$ = 8; ELS $n$ = 8 males). $n$, individual males; CPM, counts per million; DiffSPG, differentiating spermatogonia; SPG, spermatogonia; SSC, spermatogonial stem cell.
